# Neuronal Cells Confinement by Micropatterned Cluster-Assembled Dots with Mechanotransductive Nanotopography

**DOI:** 10.1101/347245

**Authors:** Carsten Schulte, Jacopo Lamanna, Andrea Stefano Moro, Claudio Piazzoni, Francesca Borghi, Matteo Chighizola, Serena Ortoleva, Gabriella Racchetti, Cristina Lenardi, Alessandro Podestà, Antonio Malgaroli, Paolo Milani

## Abstract

The *in vitro* fabrication of neural networks able to simulate brain circuits and to maintain their native connectivity is of strategic importance to gain a deep understanding of neural circuit physiology and brain natural computational algorithm(s). This would also enable a wide-range of applications including the development of efficient brain-on-chip devices or brain-computer interfaces. Chemical and mechanotransductive cues cooperate to promote proper development and functioning of neural networks. Since the 80’s, controlled growth of mammalian neuronal cells on micrometric patterned chemical cues with the development of synaptic connections and electrical activity has been reported, however the role of mechanotransductive signaling on the growth/organization of neural networks has not been investigated so far. Here we report the fabrication and characterization of patterned substrates for neuronal culture with a controlled structure both at the nano- and microscale suitable for the selective adhesion of neuronal cells. Nanostructured micrometric dots were patterned on passivated cell-repellent glass substrates by supersonic cluster beam deposition of zirconia nanoparticles through stencil masks. Cluster-assembled nanostructured zirconia surfaces are characterized by nanotopographical features that can direct the maturation of neural networks by mechanotransductive signaling. Our approach produces a controlled microscale pattern of adhesive areas with predetermined nanoscale morphology. We have validated these micropatterned substrates using a neuronal cell line (PC12 cells) and cultured hippocampal neurons. While cells have been uniformly plated on the substrates, they adhered only on the nanostructured zirconia regions, remaining effectively confined inside the nanostructured dots on which they were found to grow, move and differentiate.

## INTRODUCTION

Despite the staggering complexity of the brain and its connectome, the nervous system computes and controls behaviors by activating subsets of neural circuits and modules characterized by a simple design and composed of few subtypes of neuronal cells with comparable activity patterns. A fundamental issue in neuroscience therefore relates to the questions about how such a variety of complex computational abilities emerge when these elementary networks of cells are connected using specific architectures and hierarchies of communication (1),(2),(3).

Considerable efforts have been directed to develop *in vitro* culture strategies and tools to realistically reproduce the neuron development and growth while governing the complexity of neural population organization (4),(5),(6),(7),(8). The goal is to gain a deeper understanding of and control over the interaction of single building blocks concurring to brain complexity and to disentangle the relationship between structural organization and functional behavior (4),(5),(6),(7),(8),(9),(10). The potential applications are certainly manifold and versatile, ranging from the comprehension of the network communication and crosstalk between different types of neurons, to the realization of alternatives for animal models and brain slices to study neural disorders or the development of drug testing platforms in brain-on-chip approaches (4),(5),(6),(7),(8).

In order to reproduce neural architectures observed *in vivo*, it is necessary to achieve a spatial control *in vitro* by constraining the adhesion of individual living neurons on predefined substrate areas and make them follow specific connectivity designs, in conditions where the number and subtypes of neuronal cells and their connectivity patterns are precisely determined *a priori* (4),(5),(6),(7),(8).

Different approaches have been proposed based on the patterning of chemical adhesive cues on the substrate as effective ways to restrict the adhesion of neuronal cells to defined spatial locations defined in advance and then guide their neurite outgrowth (11). Spatial control over network geometry and topology, with micrometric resolution, has been demonstrated by using lithographic techniques typical of semiconductors to define regions with selected adhesive or antifouling properties (12). To date, the leading approaches used to implement these objectives are based on the combination of cell-repelling, antifouling surfaces with chemical patterning (e.g. by microcontact printing of components of the extracellular matrix) or microstructuring by microfabrication techniques typical of microelectromechanical systems (4),(6),(7),(8).

Microfluidic devices have also demonstrated capabilities for maintaining and studying neuronal cells and circuits in stable micro-culture. By using replica molding environments approximating not only single cells, but even single neuronal processes, can be fabricated and the control on spatial and temporal cues within microdevices enables the study of neurons and their processes (6),(7),(8),(13).

A largely neglected aspect and thus unexploited potential in neural circuit engineering approaches are biophysical substrate cues (7) (such as e.g. nanotopography and rigidity (14)), despite the increasing awareness about their importance for neuron development (7),(15). The neuron/nanoenvironment interaction has an acknowledged strong impact on neuronal functioning and behavior, particularly regarding neurito- and synaptogenesis (7),(15),(16),(17),(18). Chemical cues play a critical role in developing and maintaining neuronal function, however it has recently been recognized that physical cues due to the stiffness and nanoscale topography of the extracellular matrix are also of critical importance for the maintenance and regulation of neuronal function, although this role is not yet well understood (17),(18).

Nanoscale roughness modulates the survival and function of different cellular components of the central nervous system (7),(15),(17). Neurite/axon outgrowth, synaptogenesis and network maturation are fostered by complex mechanotransductive processes and signaling triggered by the neuronal interaction with appropriate nanotopographical surface features (15),(17),(18),(19),(20),(21),(22). It would be thus very important to develop approaches for the fabrication of patterned substrates able to provide controlled nanoscale topographical cues to study the role of mechanotransduction on the formation of neuron networks.

Recently we have reported that transition metal oxide nanostructured surfaces, in particular zirconia, produced by supersonic cluster beam deposition (SCBD) have extracellular matrix-like nanotopographical features (20),(23) promoting neuronal differentiation by mechanotransductive processes and signaling (19),(20),(21),(22). Primary neuronal cells can be cultured on these cluster-assembled substrates where they display an accelerated maturation and build-up of a functional network of neurons and synapses (21). Furthermore, SCBD enables a facile scale-up of the surface area with a predefined reproducible nanotopography (23),(24), making this nanofabrication technique compatible with thorough molecular characterization of these cell networks, including omics approaches (20),(21),(22) and high-throughput applications (25).

Here we demonstrate the possibility of confining neuronal cells on micropatterns with a defined nanoscale topography fabricated by SCBD. The micropatterns have been tested and validated on a neuronal-like cell line (PC12) and on CA3-CA1 primary hippocampal neurons that grew, moved and differentiated on the adhesive nanostructured zirconia areas but scarcely overcame the boundaries between adhesive and cell-repellent territories.

## RESULTS AND DISCUSSION

### Micropattern fabrication

In previous papers we reported the fabrication of nanostructured ZrO_2_ (ns-ZrO_2_) substrates with disordered yet controlled topographical features based on the assembling of zirconia nanoparticles, produced in the gas phase and accelerated in a supersonic expansion, on flat substrates (23). This bottom-up assembling technique produces nanostructured films with a nanoscale topography accurately controlled in a reproducible manner (23),(24),(26),(27).

We deposited nanostructured zirconia (ns-ZrO_2_) micropatterns on glass substrates previously functionalized with an antifouling cell-repellent surface treatment. To render the culture substrate repulsive for cell adhesion, we used the antifouling copolymer PAcrAm-*g*-(PMOXA, NH_2_, Si) which has been optimized, compared to the current gold standard PLL-*g*-PEG (28), specifically for bottom-up neuroscience applications (29). The copolymer was deposited on a glass surface by evaporation in a vacuum desiccator. Subsequently, the patterned deposition of the cluster-assembled nanostructured zirconia film was obtained by placing a stencil mask with the wanted micropattern design (fabricated by laser cutting, examples for the apertures in Fig. S1) onto the passivated glass surface. SCBD allows obtaining a very high lateral resolution for the production of micropatterns via stencil masks and can deposit cluster-assembled films on very delicate substrates without damage or significant heating of the surface (30). We thus obtained a geometric partitioning of zones with nanotopographical features suitable for cell adhesion on an elsewhere cell-repellent surface (Fig. 1).

**Fig. 1.**
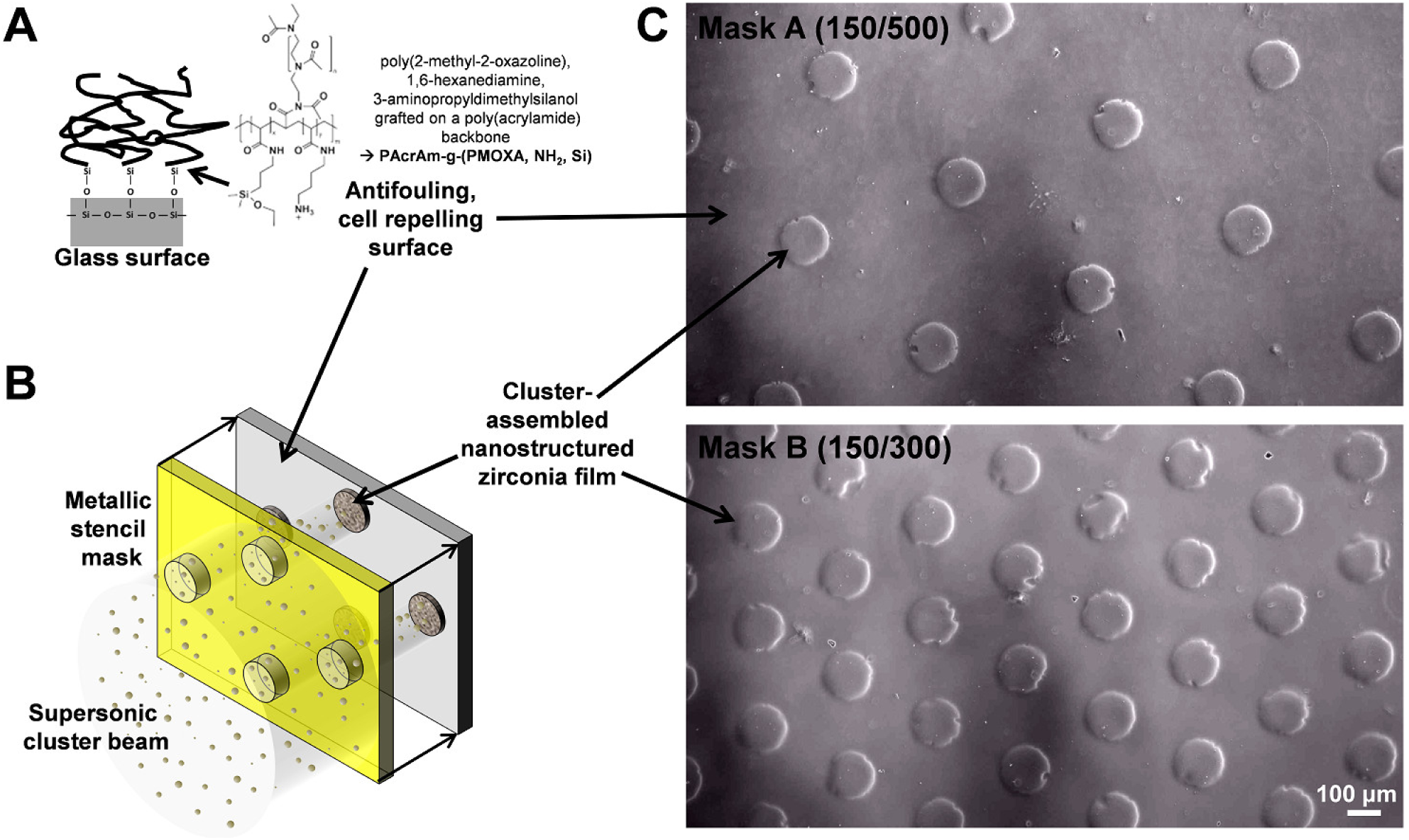
(A, B) Schematic representation of the different steps of the micropattern fabrication based on SCBD. (A) The surface of a glass slide is passivated by covalently linking the antifouling copolymer PAcrAm-*g*-(PMOXA, NH_2_, Si) to the hydroxylated glass. The chemical formula of the copolymer is a reprint with permission from Weydert et al. (29), Copyright 2018 American Chemical Society. (B) On these passivated surfaces zirconia nanoclusters are deposited through stencil masks to achieve the micropatterns with nanotopographies. (C) Representative phase contrast images of two micropatterns produced by SCBD are shown, both with circular nanostructured film dots that have diameter of 150 µm, distinguished by the center-to-center distance which is 500 µm for Mask A (150/500) and 300 µm for Mask B (150/300).

### Physical characterization of the micropatterning

Fig. 2 shows an atomic force microscopy (AFM) image of a single ns-ZrO_2_ dot, characterized by a diameter of 150 µm and a thickness of 240 nm. The typical surface morphologies of the patterned ns-ZrO_2_ (central region) and the passivated substrate areas are displayed in the upper lateral boxes of Fig. 2. Representative topographical profiles are superimposed to the central image of Fig. 2 in order to appreciate the geometrical regularity and the symmetry of the ns-ZrO_2_ dots. In particular, the dot possesses a large central plateau with root mean square (rms) roughness R_q_ = 14.7 ± 0.5 nm. The ns-ZrO_2_ film forming the dots in the micropattern is characterized by the assembly of nanoscale building blocks forming a rough layer, with high specific-area and porosity at the nano- and sub-nanometer scale (24),(27),(31), as clearly visible in the top-left box of Fig. 2.

**Fig. 2.**
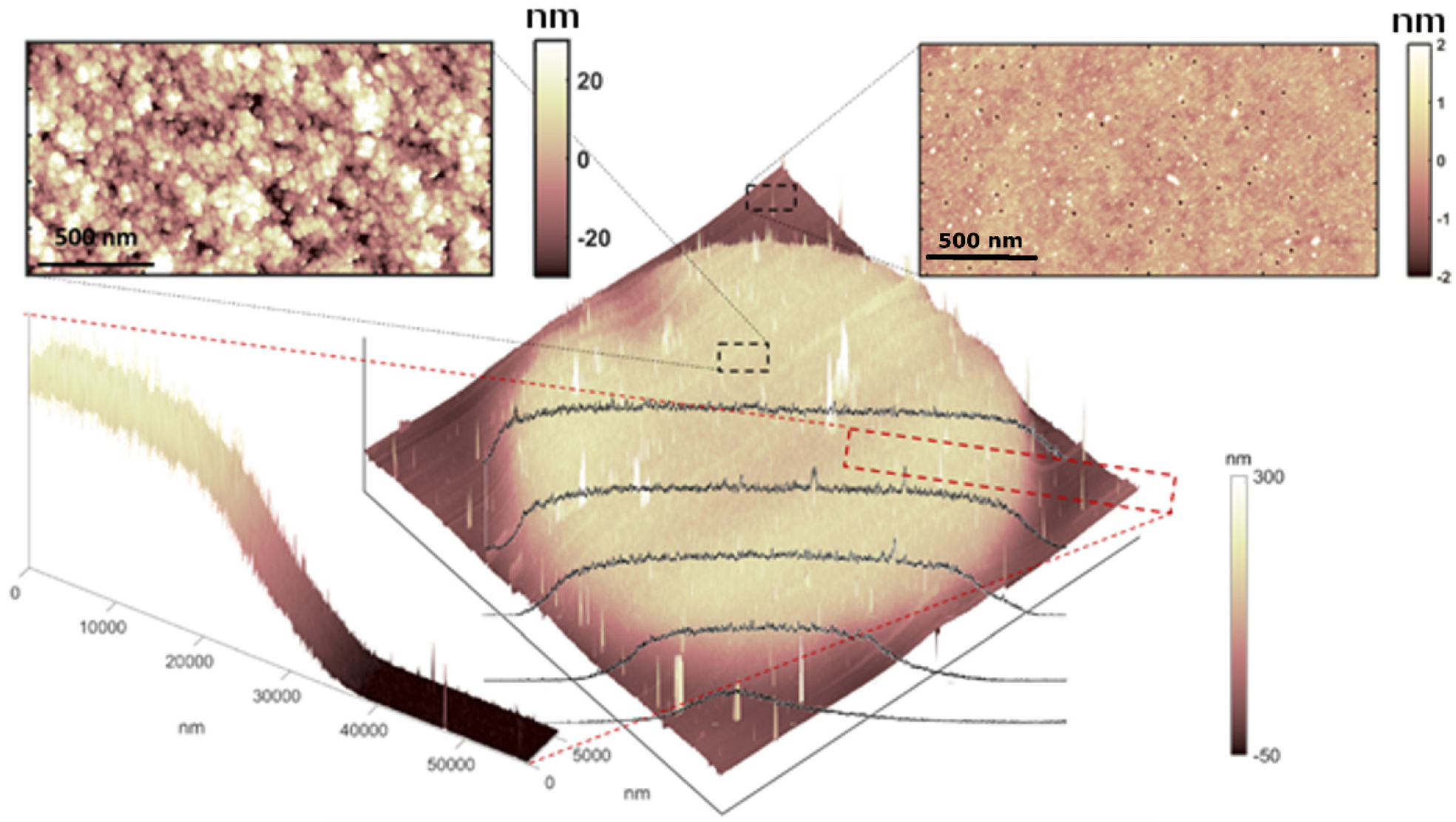
Representative 3D AFM map of a patterned ns-ZrO_2_ dot, and the adjacent passivated surface (image dimension 150 µm x 150 µm x 0.35 µm). The upper lateral boxes show in detail the surface morphology of the ns-ZrO_2_ film (left), and of the passivation layer (right). The bottom-left inset shows a 3D profile map recorded across the dot edge (same vertical scale of the central map).

The top-right box of Fig. 2 shows that the passivation layer is molecularly thin, with a roughness of only 0.3 ± 0.1 nm (similar to the roughness of the clean substrate, which is 0.1 nm; the small nanometer-sized holes typical of the glass coverslip surface can be clearly seen). A narrow three-dimensional topographical map recorded at the dot edge is shown in the bottom-left corner of Fig. 2, demonstrating that the roughness increases from 3 nm at the bottom up to 15 nm on the upper plateau, across a 30 µm-wide thickness gradient.

The morphological properties of the cluster-assembled film evolve with deposition time and film thickness. The rms roughness increases, according to a simple scaling law, R_q_ ~ h^β^, where β is the growth exponent (32). Fig. 3 shows the scaling of R_q_ (measured from AFM images of the central region of the ns-ZrO_2_ dots) with the increasing film thickness h, at room temperature. The characterization of the scaling exponents allows the prediction and control of the morphological properties of the nanostructured materials, in this case ns-ZrO_2_, for which we found β = 0.31 ± 0.09. This value is typical of interfaces evolving in the ballistic deposition regime (31),(32), where incoming particles stick on the substrate upon landing, with negligible surface diffusion (27).

**Fig. 3.**
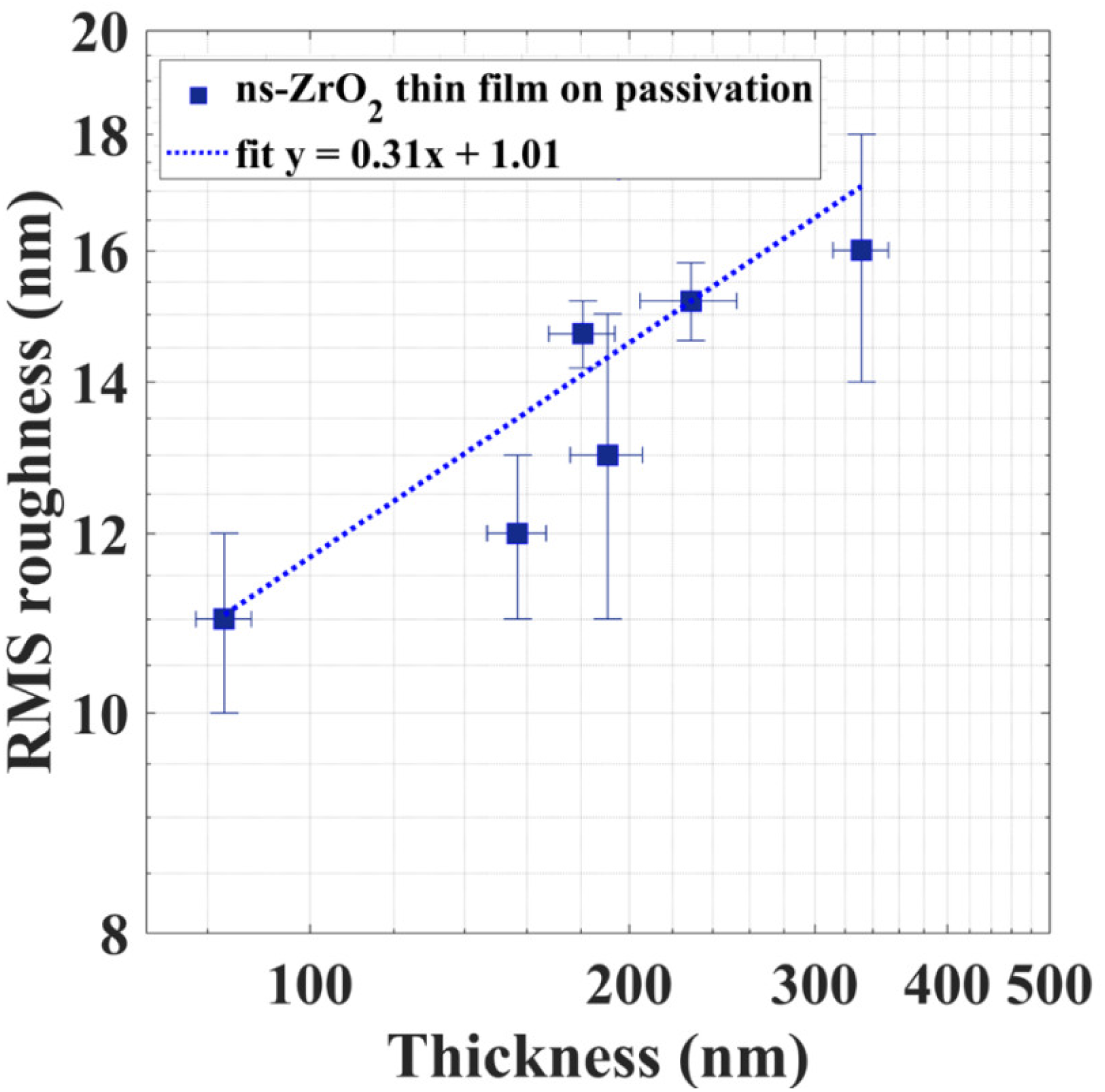
Evolution of the rms roughness R_q_ with film thickness h, for as-deposited ns-ZrO_2_ dots on passivated coverslips, in log-log scale.

Very similar values for the growth exponent have been obtained in previous studies for cluster-assembled zirconia (23),(24), and titania (31), nanostructured films produced by SCBD, deposited directly on clean oxidized silicon substrates. The thickness values measured by AFM on the patterned samples have always been found in good agreement with the values measured on reference samples co-deposited on silicon substrates (h = 240 ± 20 nm).

The deposited pattern turned out to be stable in the canonical cell culture medium over several days, and in particular the overall surface morphology was conserved (Table S1). Moreover, the concentration of residual zirconium released by the nanostructured film in the medium resulted to be lower than the limit of detection (Fig. S2).

### Neuronal cells behavior on the micropatterned substrates

We analyzed the adhesion of a commonly used neuronal cell line, the PC12 cells (Fig. 4), plated on a typical micropattern of ns-ZrO_2_ dots (Fig. 1). Cells were seeded at 3 different concentrations (ranging from 5.000 to 50.000 cells/cm^2^; Fig. 4).

**Fig. 4.**
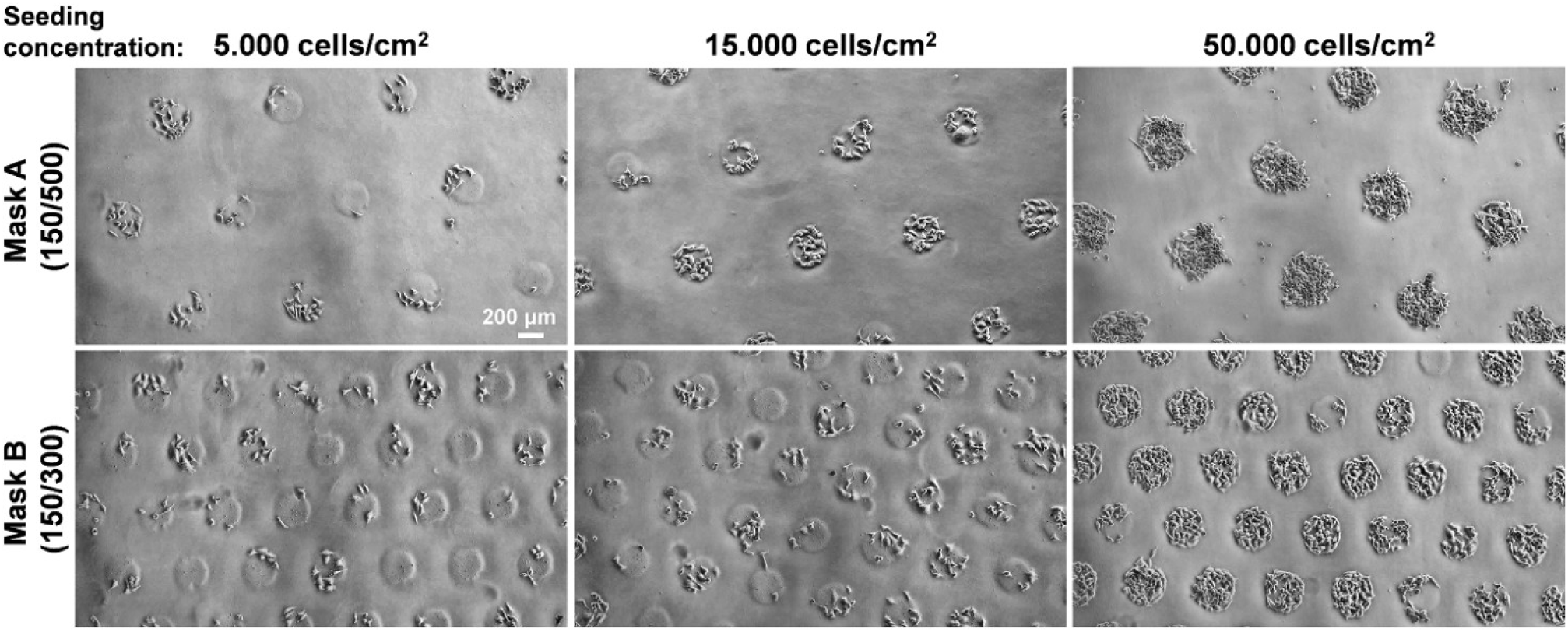
Representative phase contrast images of PC12 cells plated at the indicated concentrations on two different fabricated micropatterns. Images were taken 24 hours after cell seeding.

In these experiments we quantified the number of cells and their distribution inside and outside the adhesive ns-ZrO_2_ zones (Fig. 5). In all conditions, cells adhered predominantly to the nanostructures dots, with increasing density on the dots when the seeding concentration was augmented (Fig. 4 and Fig. 5C).

**Fig. 5.**
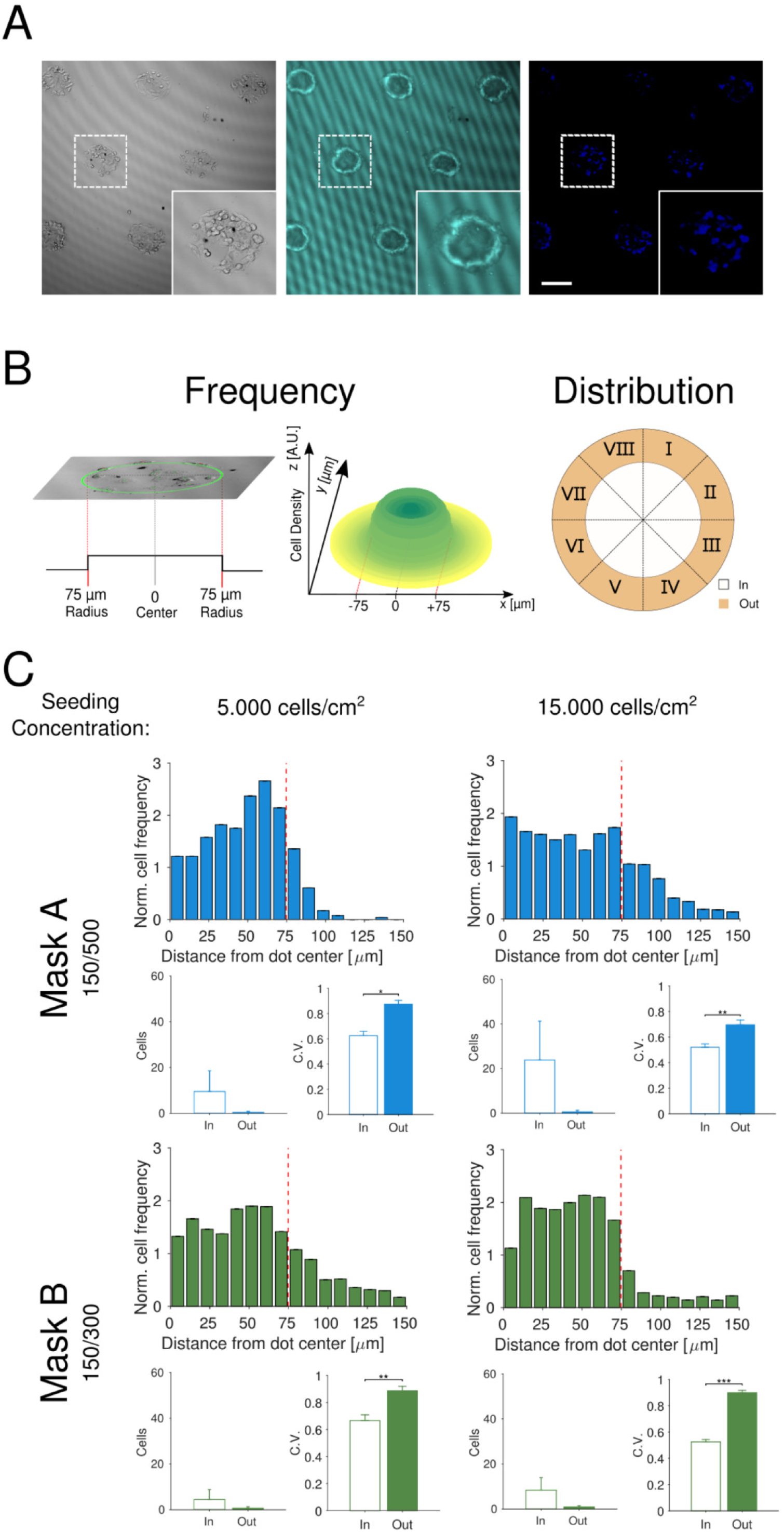
Analysis of cell density and distribution on nanostructured dots. (A) Phase contrast image of PC12 cells (left) plated for 24 hours, the same region imaged using confocal reflection microscopy to highlight the ns-ZrO_2_ dots (middle), and staining for cell nuclei (right; SYTO82 dye). Scale bar: 100 µm (B) Left: cartoon depicting the frame of reference for dot area identification and cell density estimation and the 3D representation of the obtained PC12 cell density inside and around the dot center. Right: the segmentation scheme adopted for the coefficient of variation (C.V.) analysis, where each analyzed region comprising the dot (in) and its periphery (out) is divided into 8 sub-regions. (C) The upper histograms show, for each mask (A, B), cell frequency measured in concentric bins along a 150 µm radius (bin-width 9.375 µm) starting from the center of the ns-ZrO_2_ dots, for the seeding concentrations of 5.000 (leftmost panels) and 15.000 (rightmost) cells/cm^2^. The dashed red line at 75 µm indicates dot boundary. Under each histogram, two bar-plots show the absolute average cell number obtained over (in) and outside (out) the dot, as well as the C.V. obtained inside and outside the ns-ZrO_2_ dots (values are mean ± standard deviation, stdev).

On both patterns, after 24 hours only a minor fraction of cells (4.3% - 16.2% of total cell number) could be found outside the nanostructured areas (Fig. 5C), most of these cells (2.4% - 14.1% of total cell number) being in close proximity to the dots and/or attached to other cells on them (Fig. 5C). At the highest concentration many dots were found to be overcrowded with cells, especially in the mask A (150/500) condition which provides a lower absolute adhesive area (see the corresponding phase contrast images in Fig. 4), leading to an increased cell number outside the specific adhesive zone. However, also in this case, most of the cells found outside were still in proximity to the cells on the adhesion zones. In all conditions very few cells (0.9% - 4.8% of the total cell number, see also the results of Fig. 8) actually adhered on the passivated surface area outside of the adhesive zones without any contact to the cell colonies on the dots (Fig. 5C).

The coefficient of variation (C.V.) computed for the number of cells among the different regions of ns-ZrO_2_ dots and neighboring areas (each dot/neighboring area divided into 8 equal regions, see Fig. 5A-C) showed a more homogeneous distribution of cells on the adhesive nanostructured zones than on the passivated neighboring surface outside. The higher C.V. outside is most probably due to cell aggregation on the passivated surface area because of weak adhesion, compared to the ns-ZrO_2_ areas that favor adhesion (Fig. 5C).

A morphological analysis of the cells (for the seeding concentration of 15.000 cells/cm^2^) was performed and demonstrated a lower area and perimeter of the cells outside the adhesive zones which indicates lower spreading outside, a likely consequence of impaired adhesion (Fig S3).

We performed a set of 24 hours time-lapse recordings ((Movie S1) to monitor the adhesion process starting 30 minutes after seeding (seeding concentration 15.000 cells/cm^2^). From the analysis of these recordings, a progressive increase of the cell density on the adhesive zones over time can be observed (Fig. 6A and B). In many instances, once cells encounter the adhesive zones on their trajectory (typical examples of trajectories are displayed, Fig. 6C), they adhere to it and start differentiating (examples can be seen in the (Movie S2).

**Fig. 6.**
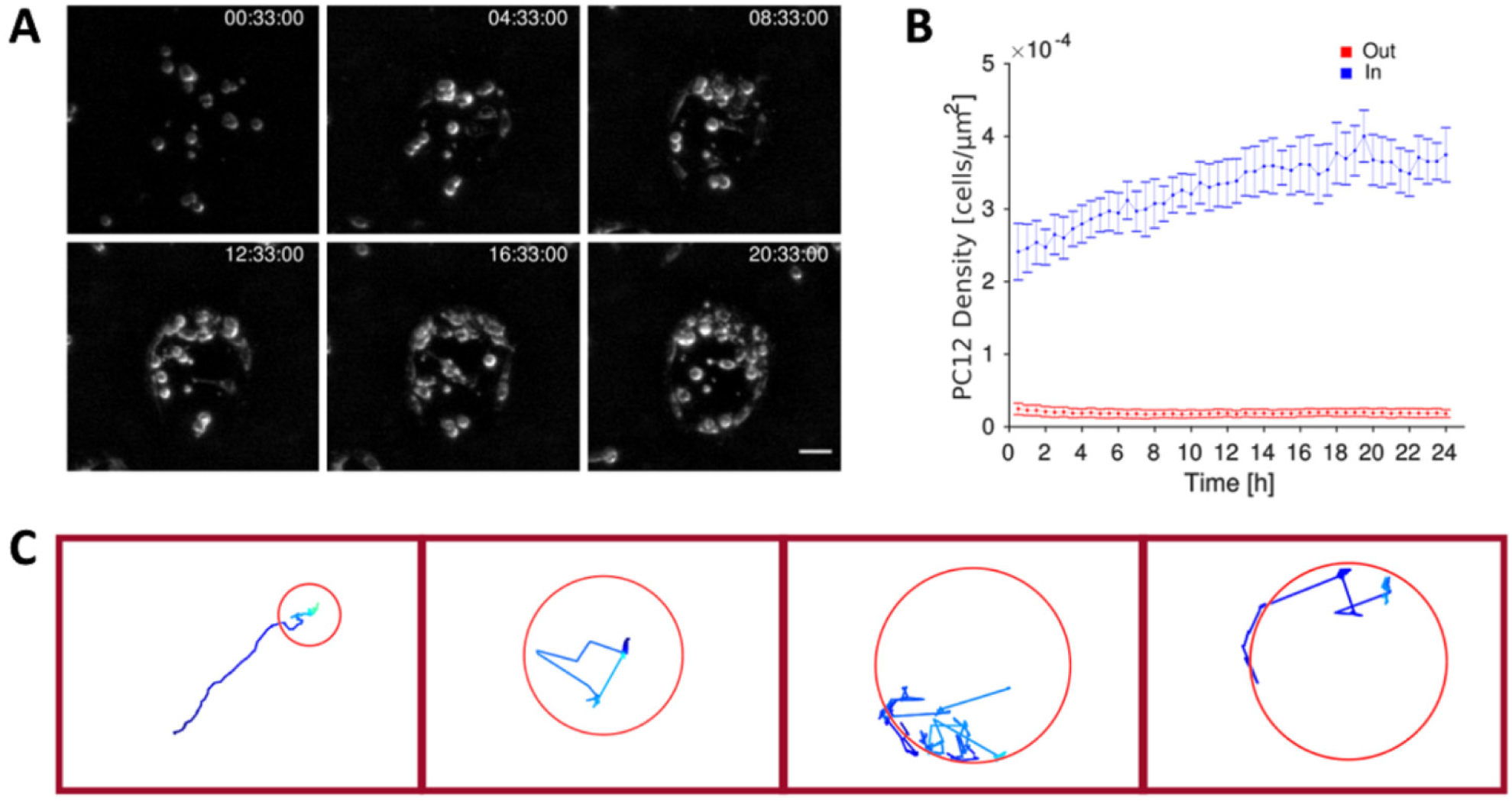
Time-lapse analysis of cell movements after plating. (A) Frames from different time points of a time-lapse recording (from (Movie S1) showing the colonization of a representative ns-ZrO_2_ dot; (B) Quantification of the time dependent evolution of cell density inside (blue dots) and outside (red dots) the nanostructured dots. (C) Examples of some representative cell trajectories.

In addition, we carried out a combined AFM and optical microscopy characterization focusing on the peripheral regions of the ns-ZrO_2_ dots, where cells find a gradient in the interfacial physicochemical properties, passing from the smooth passivated glass substrate to the rough ns-ZrO_2_ film. The inset of Fig. 7 shows a panoramic view of a ns-ZrO_2_ dot populated by the cells. The cells typically remain on the nanostructured film, minimizing, if not completely avoiding, the contact with the passivated glass (a behavior that can also be observed in (Movie S2). AFM allowed getting a deeper insight on the boundary region of the dots. Fig. 7A-E show that the cells manage to follow with nanoscale precision the curvature of the ns-ZrO_2_ dot edge, staying on the nanostructured film side, away from the passivated area.

**Fig. 7.**
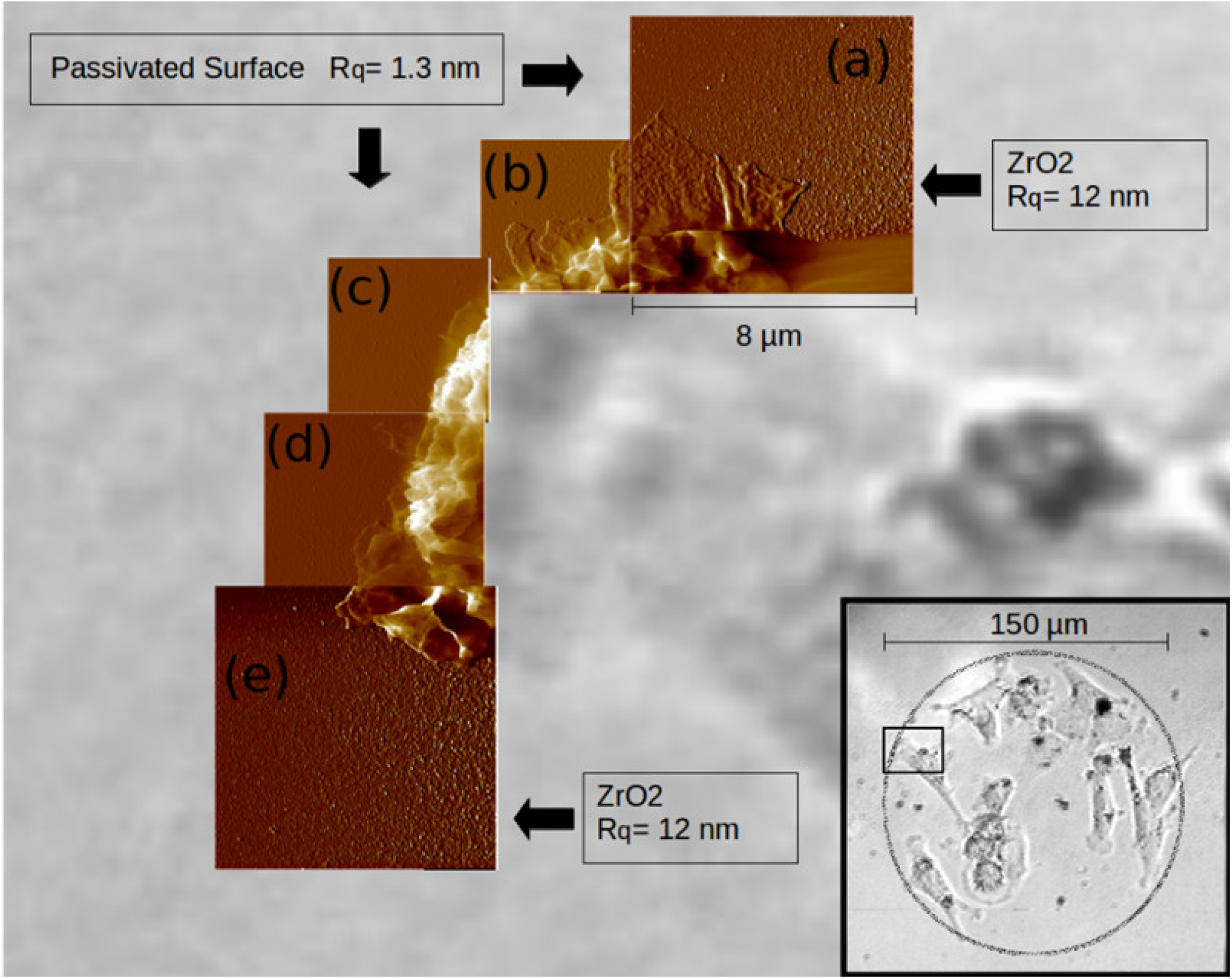
Optical image (170 μm x 170 μm) of a region of a ns-ZrO_2_ dot populated by cells (bottom-right inset) after 1 day in culture, with superimposed AFM deflection error maps.

In particular, the cells explore the whole surface of the ns-ZrO_2_ dot irrespective of the local value of the surface roughness (ranging from 3 nm at the periphery, to 15 nm in the center), avoiding the passivated substrate. The cells protrude lamellipodia, which possess a higher membrane roughness (R_q_~10 nm) when located over the ns-ZrO_2_ film, than on the passivated substrate (R_q_~2nm). The observed difference is likely due to the higher density of adhesion spots at the cell outer membrane in correspondence of the nanostructured substrate (33).

### Micropattern compliance and cell resistance of the passivated surface

An important and still challenging issue with respect to the realization of reliable and controllable *in vitro* neural networks, is the compliance of the geometric substrate partitioning and cell resistance of the surface passivation over longer periods of time (8),(29). As aforementioned the antifouling agent chosen by us was PAcrAm-*g*-(PMOXA, NH_2_, Si), a recently developed copolymer elaborated specifically for the high demands of neural pattern approaches (29).

To assess the long-term stability of the surface passivation in our micropatterning approach, we seeded the PC12 cells at a low concentration (5.000 cells/cm^2^) on the micropattern produced with mask A (150/500) and cultured them in the presence of low serum medium (1% horse serum, change of medium every day) to decelerate cell proliferation. Under these conditions, after adhering the PC12 cells grown on standard cell culture plastic covered on day 1 16.3% ±3.5 of the surface area and proliferated the next days until reaching confluence after one week (coverage 92.5% ±4.5) (Fig. 8). On the passivated only glass surfaces instead, only very few cells succeeded to adhere (at day 1, 0.9% ±0.2 coverage of the total surface area) and those cells that did attach were not able to establish efficient cell proliferation further on. In this condition, the cell surface coverage stayed over the whole time period in the range observed at day 1. On the micropattern the cells adhered, as shown before, basically only to the zones with the nanostructured film. On these adhesive dots the cells proliferated until becoming overly confluent around day 6 or 7 which led to a progressive rounding up and eventual increasing cell detachment from these dots. The surface passivation outside the adhesive nanostructured zones remained instead largely stable over time, even on day 7 only 3.9% ±2.9 of the passivated surface area was covered by cells. In addition, most of the cells growing outside remained still attached to the other cells on the dots.

**Fig. 8.**
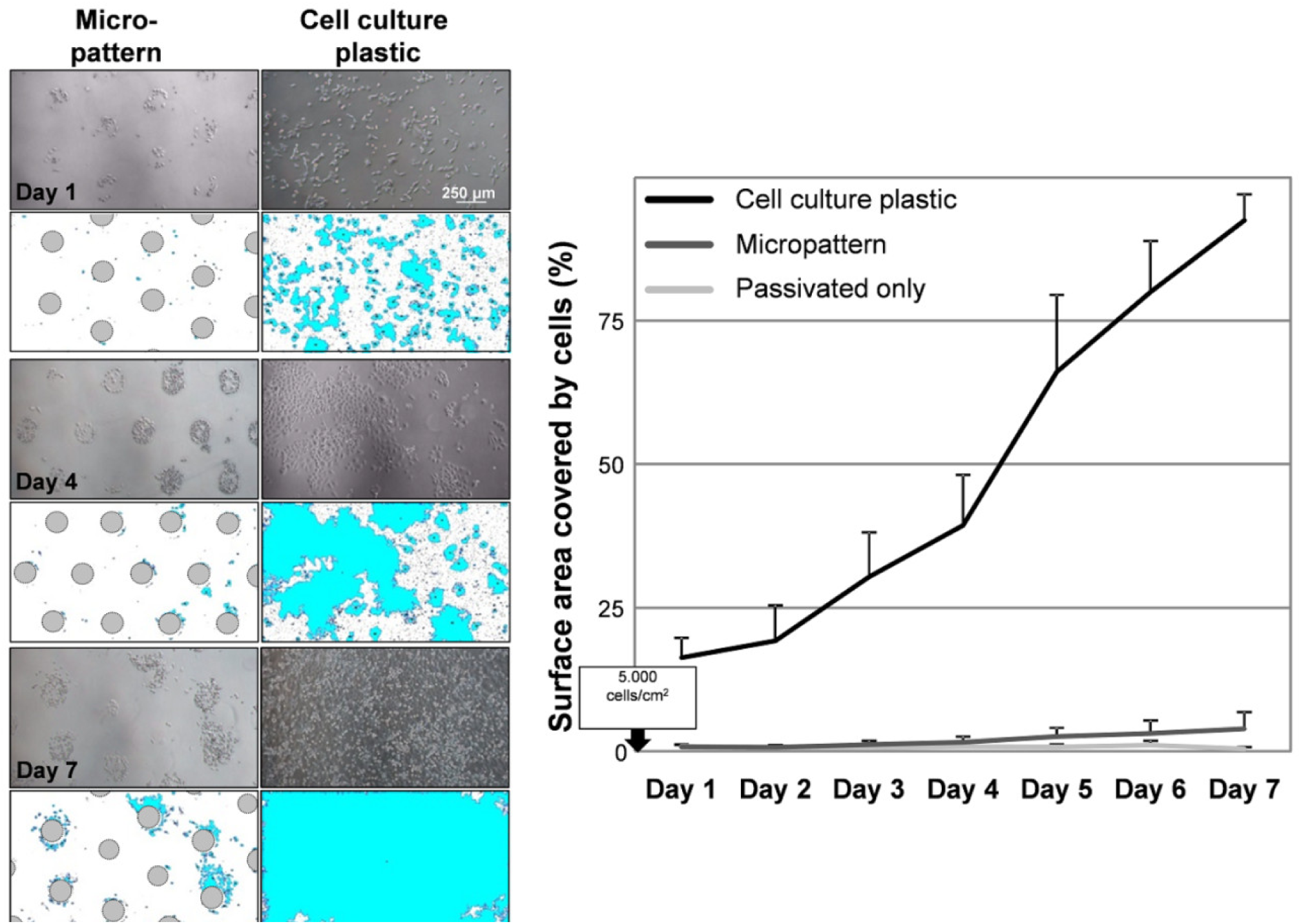
On the left, examples of phase contrast images of PC12 cells on the Mask A (150/500) micropattern or cell culture plastic at different days after plating (seeding concentration: 5.000 cells/cm^2^) are shown and underneath the corresponding images (after ImageJ processing, see Methods for details) indicating the cell coverage (in light blue) of the areas of interest (for the micropattern the passivated area and for the cell culture plastic the whole surface). The graph (lines display the average flanked with stdev) on the right summarizes the data of the analyzed conditions cell culture plastic, passivated glass only and the micropattern.

**Fig. 9.**
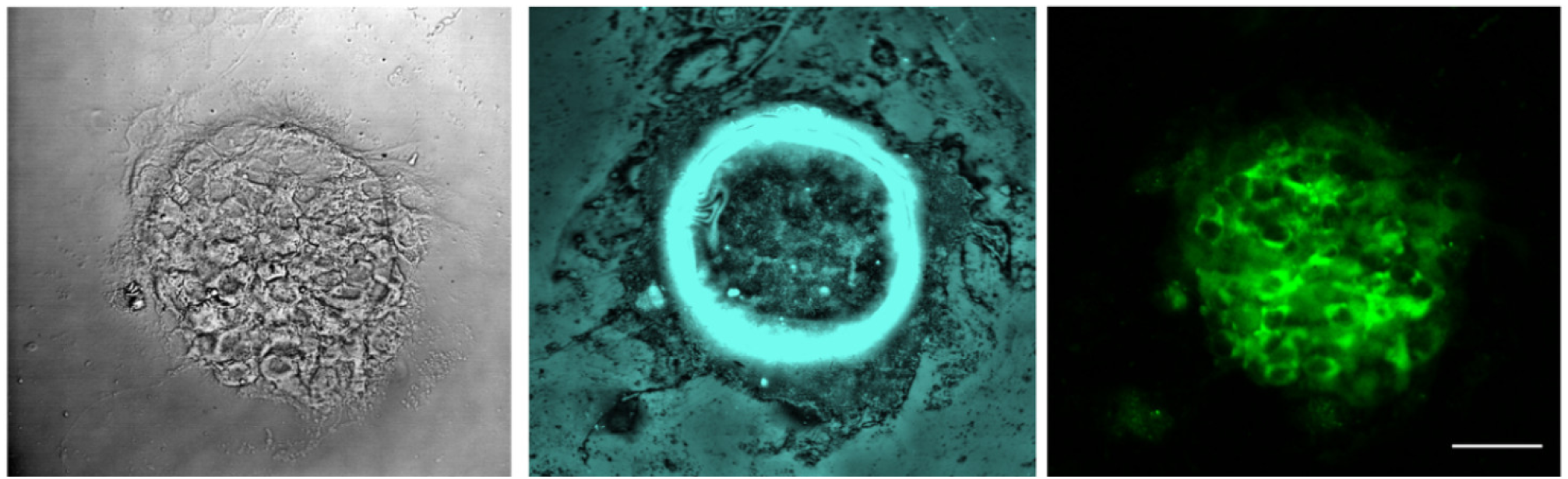
Representative images of primary rat hippocampal cells cultured on circular ns-ZrO_2_ dots. These micropatterns were fabricated with the procedure outlined in this work (left: phase contrast, middle: confocal reflection, right: NeuN staining). The cells were fixed after 7 days in culture (see also Fig. S5).

Considering the context of these experiments, it is noteworthy to point out that the clone we used in this study was PC12 Adh (ATCC^®^ CRL-1721.1™). The “Adh” stands for “adherent”, meaning that this clone is more adherent than the widely used standard PC12 cell clone. In canonical cell culture conditions (e.g. PLL-coated glass) the cell morphology of this clone is indeed characterized by distinct spreading, the build-up of large lamellipodia (see also Fig. 7 and (Movie S2) and the strong formation of focal adhesions and stress fibers (20).

Altogether, the results demonstrate therefore a high potential of this geometric partitioning in regard to confinement of cell adhesion on the designated areas.

### Primary hippocampal neurons cultured on the micropattern

We tested the culturing of primary hippocampal rat neurons on cluster-assembled micropatterns. We have previously demonstrated that these neurons mature in a more efficient way on adequate nanotopographies produced in this manner, but these experiments were done on unrestricted whole surface depositions (21). We observe that the primary hippocampal neurons selectively adhere to the cluster-assembled dots and that they can be maintained in culture for weeks (>3 weeks), comparable to neurons grown on glass coated with poly-L-ornithine and matrigel^®^. Interestingly enough, unlike the canonical hippocampal cell culture conditions, these micropatterns were used in their original state, meaning that they have not been coated with any substance prior to the cell seeding, i.e. without either poly-L-ornithine or matrigel^®^.

## CONCLUSIONS

With this work we demonstrated a method to integrate extracellular matrix-like topographical nanostructured features into micropatterns suitable for neural network fabrication. The micropatterned nanostructured zirconia surfaces were fabricated by coupling SCBD with the use of microfabricated stencil masks. The micropatterns were printed on glass coverslips that have been passivated to reduce cell adhesion by a coating with the antifouling copolymer PAcram-g-(PMOXA, NH_2_, Si).

Micropatterns with circular dots served as a proof-of-principle for the characterization of the stability, functionality and efficiency of the protocol. They were tested with two different cell types, i.e. the neuron-like PC12 cells and rat neonatal primary hippocampal cells. For both cell types a stable confinement of cell adhesion in the destined micropatterned zones was observed.

Interestingly enough, these micropatterns can be utilized in their original state and do not require coating of any type, such as e.g. the usually applied poly-L-ornithine and matrigel^®^ in the context of primary (hippocampal) neuronal cultures. This latter aspect is highly relevant because the use of matrigel^®^ e.g. comes with a cost, literally, but also in a wider sense, due to problems related to lotto-lot variability, ill-defined composition and the possibility of xenogeneic contaminants (34),(35).

Providing a reliably reproducible and controllable geometric partitioning of the cellular environment; equipped with nanotopographical features has the additional benefits of 1) being promotive for neuronal functioning/maturation and 2) allowing the avoidance of any coating, represents a strong potential compared to to-date published micropatterns, in particular in this challenging neuronal cell culture framework (8).

Micropatterns with such a fully synthetic but highly biomimetic substrate have manifold possible applications; for the highlighted development of defined neuronal network topologies and brain-on-chip devices, but also for more general approaches involving other cell types (in particular stem cells in various differentiation scenarios) and for sophisticated mechanotransductive studies.

## MATERIALS AND METHODS

### Substrate fabrication and characterization

#### Surface passivation with PAcrAm-g-(PMOXA, NH_2_, Si)

The carriers (Ø13mm glass coverslips) have been cleaned with Piranha solution in order to remove organic residues on the glass surface and make them hydrophilic. The Piranha clean is a mixture of H_2_SO_4_:H_2_O_2_ (3:1) and, since it is a strong oxidant solution it hydroxylates glass surfaces. After preliminary cleaning the substrates are accurately washed with ultrapure water (milliQ) and dried under pure nitrogen flux.

After that the coverslips are closely covered with an antifouling coating achieved by a new copolymer PAcrAm-g-(PMOXA, NH_2_, Si) (for the full chemical name, see Fig. 1) produced by SuSoS AG (Duebendorf, Switzerland) (28),(29). The backbone of this polymer is poly(acrylamide) (PAcrAm). Several components are grafted on this backbone; the hexaneamine fosters spontaneous electrostatic adsorption in buffered aqueous solutions, the propyldimethylethoxysilane serves to augment the durability by enabling covalent bonding to the hydroxylated coverslips, and the polymer poly(2-methyl-2-oxazoline) (PMOXA) instead provides the antifouling feature (28),(29).

The coating protocol has the following steps. The freshly cleaned coverslips are placed on a custom-made (by 3D printing) support mounted inside a glass vacuum desiccator together with an Eppendorf vessel containing 10 µl of the copolymer (at a concentration of 0.1 mg/ml in HEPES, pH 7.4). The desiccator is sealed and evacuated up to the pressure of about 10^−2^ mbar and maintained at this vacuum level for 3 min in order to allow the evaporation of the high vapor pressure copolymer and the deposition on the samples. The desiccator is brought back to normal pressure and afterwards the coverslips are accurately washed several times with ultrapure water (milliQ). A thin coating of the copolymer is then formed on the exposed surface of the coverslips.

#### Micropatterning by supersonic cluster beam deposition with stencil masks

Cluster-assembled zirconia dots were deposited on the passivated substrates by a SCBD equipped with pulsed microplasma cluster source (PMCS). The details of the deposition technique and PMCS can be found elsewhere (23),(26),(36). In brief, a zirconium rod placed in the PMCS is ablated via a pulsed plasma discharge supported by the injection of Ar pulses, taking place in the source cavity faced on a high-vacuum chamber. The ablated species thermalize with the injected inert gas and condense to form zirconia clusters; the cluster-Ar mixture is then expanded through a nozzle into a vacuum to form a supersonic beam (37). The cluster beam impinges on a substrate placed on a sample holder placed in a second vacuum chamber. A controlled movement of the sample holder (rastering) allows getting uniform coverage of the substrate surface. A quartz microbalance placed in the rastering area provide an on-line monitoring of the deposition rate. Deposition process parameters are controlled in order to select specific characteristics of the resulting film and, in particular, the surface roughness (23),(31). The produced samples had a target surface roughness fixed to 15 nm, as it proved to be best-performing in regard to PC12 differentiation (20).

Patterns are obtained through the interposition of a stencil mask (26) (see Fig. 1). The masks used in this work were produced by laser machining of a 150 μm thick steel foil. Two types of patterns were used:

1. Mask A: Dots of 150 μm diameter and 500 μm center-to-center distance (150/500)
2. Mask B: Dots of 150 μm diameter and 300 μm center-to-center distance (150/300)

In both patterns dots are displaced in a hexagonal matrix fashion. Fig. S1 shows representative phase contrast images of Mask A (150/500) and Mask B (150/300), respectively. A suitable custom-made sample holder was used to host the passivated coverslips and fix the masks in contact with them.

The assessment of the samples by atomic force microscopy (AFM) and stylus profilometry guaranteed that the expected dimension of dots and center-to-center distance were carefully replicated.

#### Characterization of surface morphology

The surface morphology of ns-ZrO_2_ patterns on passivated substrates was characterized using a Multimode 8 AFM produced by Bruker. Several 2 µm × 1 µm images of the ZrO_2_ micropatterns were acquired in the native state after the deposition and after three days in the cell culture medium, in order to characterize the surface nanoscale morphology. The AFM was operated in air in Peak-force Tapping Mode, using silicon nitride cantilevers mounting single crystal silicon tips with nominal radius below 10 nm, and resonance frequency in the range 50–90 kHz, scan rate of 1 Hz, and sampling resolution 2048×512 points. The images were flattened by line-by-line subtraction of first and second order polynomials in order to remove artifacts due to sample tilt and scanner bow. From flattened AFM images, the root-mean-square surface roughness R_q_ was calculated as the standard deviation of surface heights. The thickness of ns-ZrO_2_ dots was calculated by AFM imaging across the step at the edge of the dot, and by using a stylus profilometer (KLA – Tencor P-6, Milpitas, USA). For the sake of comparison, during the depositions aimed at producing patterns, ns-ZrO_2_ films were also deposited on monocrystalline silicon substrates (5 mm x 5 mm) by partially masking the substrate in order to produce a sharp step, and subsequently their thickness was measured. The thickness values measured in the different conditions did all agree quite well.

#### Analysis of zirconium traces by inductively coupled plasma optical emission spectrometry (ICP-OES)

For the analysis of potential zirconium traces released into the cell culture medium, the cell culture medium was incubated for 3 days (at 37°C in the cell incubator) with 5 substrates, i.e. 3 different ns-ZrO_2_ micropattern, passivated glass only and nude glass (the latter two as negative controls). The collected cell culture medium samples have been freeze-dried and have then been digested in a mixture of H_2_O_2_ and HNO_3_ (volume ratio 1:5). The instrument (ICP-OES (iCAP 6200 Duo upgrade), Thermofisher, Waltham, USA) has been calibrated with a blank and two standards containing 5, respectively 10 ppb (part per billion) of zirconium at the emission wavelength of 339.198 nm (see Fig. S2).

### Cell biological experimental methods and analysys

#### PC12 cell cultures

The PC12 clone used for this work was PC-12 Adh (ATCC Catalog no. CRL-1721.1™). The cells were maintained in RPMI-1640 medium supplemented with 10% horse serum, 5% fetal bovine serum, 2 mM L-Glutamine, 10 mM HEPES, 100 units/ml penicillin, 1 mM pyruvic acid, and 100 μg/ml streptomycin (all reagents from Sigma-Aldrich, Saint Louis, USA, if not stated otherwise) in an incubator (Galaxy S, RS Biotech, Irvine, UK) at 37°C and 5% CO_2_ (98% air-humidified). Routine subculturing was done every 2-3 days by detaching the cells with a trypsin/EDTA solution and, after centrifugation (1000x g, 5 min), resuspending the pelleted cells in an appropriate dilution.

For the experiments the cell detachment was executed with 1 mM EDTA in HBSS. After centrifugation at 1000x g for 5 min and resuspending the pellet, cell counting was performed with an improved Neubauer chamber. The cells were subsequently diluted according to the indicated cell concentrations for the individual experiments and analyses, in the full serum cell culture medium (as above) for the experiments presented in Fig. 4-7, or, for the micropattern compliance experiment (Fig. 8) in low serum medium (RPMI-1640 with 1% HS, 2 mM L-Glutamine, 10 mM HEPES, 100 units/ml penicillin, 1 mM pyruvic acid, and 100 μg/ml streptomycin). The phase contrast images of Fig. 1, 4, 8, and S1 were taken with an Axiovert 40 CFL microscope (Zeiss, Oberkochen, Germany) equipped with 20x/0.3 ph1, CP-ACHROMAT 10x/0.25 Ph1, 5x/0.12 CP-ACHROMAT objective and with a HD photo camera (True Chrome HD II, TiEsselab) operated by ISCapture imaging software.

#### Rat hippocampal cultured cells

The neonatal primary rat hippocampal cells were isolated and prepared for culturing as described previously (21),(38), with the difference that the substrates, i.e. micropatterns, were not coated with poly-L-ornithine or matrigel^®^ prior to the cell plating. All the procedures were executed according to the research and animal care procedures approved by the institutional animal care and use committee for good animal experimentation of the Scientific Institute San Raffaele complying with the code of practice for the care and use of animals for scientific purposes of the Italian Ministero della Salute (Ministry of Health) (IACUC number: 728).

#### SynaptoZip transfection

For eGFP-SynaptoZip (39) expression, neurons were transduced with GZ-carrying lentiviral vectors prediluted in culture medium 1h after plating (final titer *in vitro*: ~10^6^ TU ml-1; viruses preparation as in Ferro et al. (2017) (39).

#### Measurement and analysis of cell distribution and morphology

The fixed cells (paraformaldehyde 4% for 10 min) in the different indicated experimental conditions were washed with PBS and the nucleus was stained with SYTO82 (4 μM, 30 min, RT). The samples were mounted (FluorSave™, Merck Millipore, Burlington, USA) and the images were recorded with a confocal microscope (Leica TCS SP5, Leica, Wetzlar, Germany). The confocal reflection microscopic technique (CRM) was employed to visualize the zones with the ns-ZrO_2_ film (as torus-shaped geometry, see e.g. Fig. 5A). On the basis of this visualization we calculated their centroids and created a virtual radius of 75 μm. In addition, the maximum possible circular area around each ns-ZrO_2_ dot was considered that does not lead to superimposition with the nearest dots, valid for both patterns, which corresponds to a 300 μm diameter.

This information was used to quantify the cell frequency (cells were identified by the nuclear SYTO82 staining, see e.g. Fig. 5A) within the dot and outside along a 150 μm radius (with 16 binning steps of 9.3750 μm, see Fig. 5B). MATLAB software (MathWorks, Natick, USA) with an adapted contour algorithm published by Kong et al. (2015)(40) was utilized for the quantification. The frequency was eventually normalized with respect to the total average.

For the analysis of the coefficient of variance (CV, i.e. standard deviation/mean) a ring around the ns-ZrO_2_ dots was considered with an area that equals the area of the ns-ZrO_2_ dots. Each dot and ring area was segmented into 8 equal regions (see Fig. 5B for visual representation) and a CV index for the number of cells inside versus outside was calculated.

The comparison of the absolute cell numbers inside and outside the dots (for the latter only cells in vicinity of the dots still in contact with cells on the dot were considered) were determined by manual counting from the phase contrast images with the help of ImageJ (NIH, New York, USA). We decided for this manual approach to obtain the absolute cell numbers because the absolute numbers were generally underestimated by the abovementioned algorithm in the conditions with high cell seeding concentrations due to overcrowding which led to difficulties in detecting and distinguishing single nuclei. Also cells on the passivated areas without any contact to cells on the dots or in their vicinity were counted separately in this way; the percentages of these cells in relation to the total cells outside are reported in the main text.

All these parameters, i.e. cell frequency, C.V. and average cell numbers, were reported for the seeding concentrations 5.000 and 15.000 cells/cm^2^ in Fig. 5. For the seeding concentration of 50.000 cells/cm^2^ the cell frequency and C.V. could not be determined accurately due to the overcrowding of the cells on the dots which compromised algorithm reliability in distinguishing single nuclei. However, the graphs of the average cell number in and outside the dot area for this condition are reported in Fig. S4.

The graphs in the figures display the global statistic of two independent experiments; for the cell frequency in total 29 - 120 dots and for the absolute cell numbers 1043 - 2147 dots were quantified for the analyses.

An analysis of cell morphological features was performed for the experimental condition with a concentration of 15.000 cells/cm^2^ plated on the Mask B (150/300) pattern. The phase contrast images served to reconstruct the 2D membrane shape (see Fig. S3) with the help of MATLAB software by adapting a phase contrast image processing published by Jaccard et al. (41) and the abovementioned contour algorithm (40). By these means the cell area and perimeter were quantified.

#### Time-lapse imaging

The time-lapse was performed with an inverted microscope (Axiovert 200, Zeiss) equipped with a custom-made chamber to hold the sample and to secure constant culture conditions. The conditions were kept at 35°C by temperature control (Tmpcontrol 37-2 digital (2 channel), PeCon GmbH, Erbach, Germany) and 100% humidity. The cells were kept in full serum medium (see above) which was supplemented with 25 mM HEPES to stabilize the pH. Phase contrast images were taken every 3 min at very low light intensity and long exposure time to reduce phototoxicity of HEPES and cell stress. The time-lapse analysis was done with MATLAB software (MathWorks) using an adapted contour algorithm (40) for segmentation and a tracking algorithm to study the trajectories of the cell movement.

#### Micropattern compliance and cell coverage analysis

The analysis of the compliance of the micropattern and passivated cell-repellent area was performed by plating PC12 cells with a concentration of 5.000 cells/cm^2^ onto canonical cell culture plastic, passivated glass only or on the mask A (150/500) micropattern in low serum (1% HS) cell culture medium (which was changed every day). Phase contrast images of the cells in the different conditions were recorded every day for 7 days and evaluated by ImageJ. The phase contrast images were converted into 8bit images and the “Find Edges” function was applied to increase the contrast between cells and background. For the cell culture plastic and passivated only conditions these images were the basis of the threshold implementation and quantification. For the images with the micropattern, in addition the contours of the nanostructured film zones present in the image were marked and their area was measured. Afterwards these dot areas were cleared out of the image (by setting their pixel intensity to zero) to not be further considered for the subsequent quantification. Thresholds were set and applied in order to eliminate the background (i.e. the surface area not covered with cells, see white area in representative images in Fig. 8) and the “Analyze particle” function was used to detect the area covered by the cells (a particle size threshold of 50 μm^2^ (to infinity) was put to exclude unspecific debris or dead cells floating in the PBS during image recording, “include holes” was marked and no restrictions were set in regard to circularity), see light blue areas as examples in the representative images in Fig. 8. The total area of these particles for the whole image was summed and set in relation to the whole surface area (for the conditions: cell culture plastic and passivation only), or the whole surface area minus the area covered by the nanostructured film zones (marked and determined before) respectively.

The graph in Fig. 8 represents the global statistic of two independent experiments with in total 12 – 18 quantified fields.

#### Imaging of cells by AFM

Cells were fixed with 4% paraformaldehyde (10 min) after one day spent on the micropattern. The cell morphology was characterized using a Bioscope Catalyst AFM from Bruker. The AFM images (with scan size of 8-6-4 µm) were acquired in PBS buffer in Peak-Force Tapping Mode. Due to the significant height variations within the samples (up to several microns), short cantilevers with tip height of 14 µm were used (VistaProbes CS-10). The scan rate was set to 0.6 Hz, and the sampling resolution to 4096×1024 points. The accurate alignment of the optical and the AFM images were obtained using the Miro module integrated in the AFM software

## SUPPLEMENTARY MATERIAL

See supplementary material for five supplementary figures (Fig. S1 to S5), one table (Table S1), two movies (movies S1 and S2), and additional discussion.

## ACKNOWLEDGEMENTS

PM and CS acknowledge support from the European Union project “FutureNanoNeeds” grant “Framework to respond to regulatory needs of future nanomaterials and markets” (FP7-NMP-2013-LARGE-7). AP thanks the University of Milan for financial support under the project “Transition Grant 2015/2017 – Horizon 2020”. We would like to thank Federico Pezzotta from the Mechanical Workshop of the Department of Physics at the University of Milan for the help with the production of the custom-made sample holders. We thank Simona Argentiere for the help with the zirconium trace experiment.

## AUTHOR CONTRIBUTIONS

The project conceptualization was realized by AP, JL, CL, AM, PM, and CS. The principal part of the manuscript was written by ASM, JL, AM, PM and CS. FB, MC, CP, SO, CL and AP contributed to the paragraphs concerning the micropattern fabrication and the physical characterization of the micropattern, as well as the AFM investigation of the cells on the micropattern. The surface passivation and micropattern fabrication was done by SO, CS and CP, and the physical substrate characterization was performed by MC, FB, AP and CP. The cell biological experiments and the corresponding data analyses were done by ASM, JL, GR, SO and <u>CS. SA</u> performed the zirconium trace experiment.

## ETHICS APPROVAL

All the procedures were executed according to the research and animal care procedures approved by the institutional animal care and use committee for good animal experimentation of the Scientific Institute San Raffaele complying with the code of practice for the care and use of animals for scientific purposes of the Italian Ministero della Salute (Ministry of Health) (IACUC number: 728).

## SUPPLEMENTARY MATERIAL

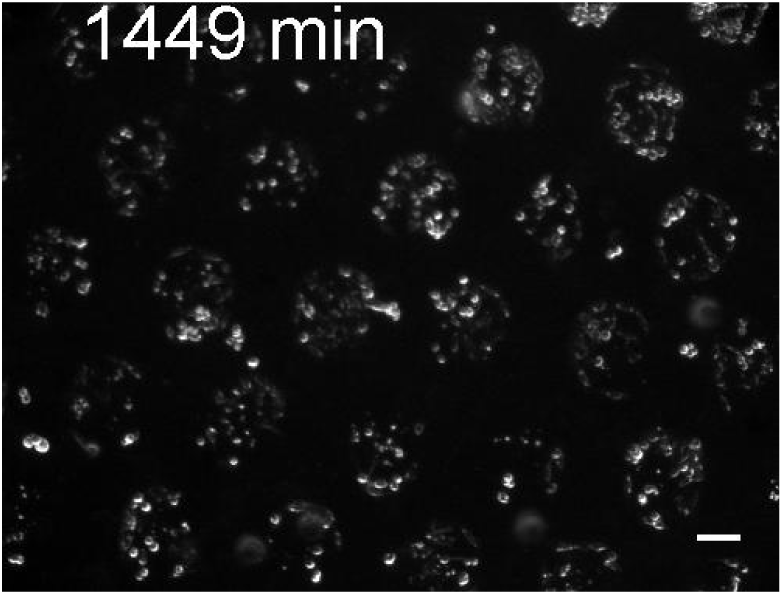
Movie S1. The movie shows a recording of PC12 cells seeded with a concentration of 15.000 cells/cm^2^ on the micropattern produced with Mask B (150/300). Scale bar = 100 μm.

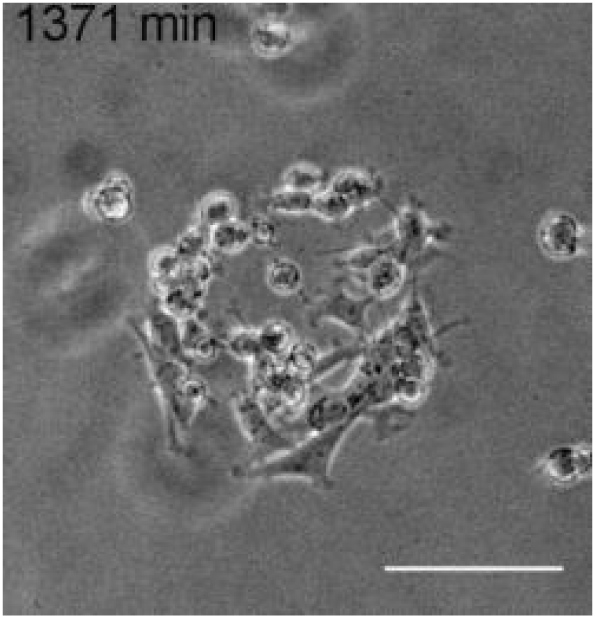
Movie S2. The movie demonstrates in detail the adhesion process of PC12 on a single ns-ZrO_2_ dot and outgrowth of neurites from several cells. Scale bar = 100 μm.

**Fig. S1.**
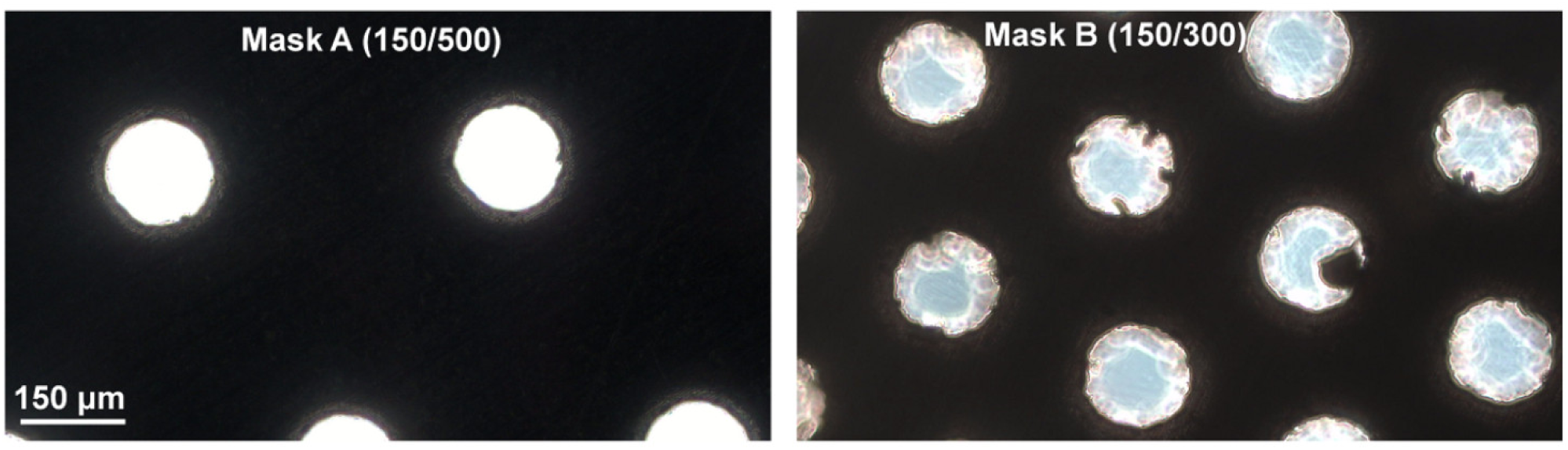
Phase contrast images of details of the metal masks used for the masked SCBD showing examples of the apertures produced by laser cutting through which passes the microscope light.

### Stability of the nanostructured pattern in the cell culture medium

We have checked the stability of the deposited material after three days in the canonical PC12 cell culture medium, i.e. RPMI supplemented with 10% horse serum and 5% fetal bovine serum. In Table I the values of the surface roughness of ns-ZrO_2_ and the passivated glass are reported. Roughness measurements have been carried out from images acquired in air, after the sample was gently washed and dried. The increase of roughness for the passivated smooth surface is attributable primarily to the deposition of proteins from the culture medium. This was confirmed by the fact that after incubation of the sample with trypsin (which digests proteins) the pristine roughness values were recovered. The same protein coating had a negligible effect on the surface morphology (and roughness) of ns-ZrO_2_. SUPPLEMENTAL

**Table I:** RMS roughness of nanostructured ZrO_2_ dots and passivated areas before and after incubation with cell culture medium, followed afterwards by trypsin treatment.

| Surface | Condition | Roughness (nm) |
| --- | --- | --- |
| ns-ZrO<sub>2</sub> dots | As Deposited | 14.7 ± 0.5 |
|  | After 3 days in culture medium | 15.2 ± 0.6 |
| Passivated area | As Deposited | 0.3 ± 0.1 |
|  | After 3 days in culture medium | 0.7 ± 0.1 |
|  | After Trypsin | 0.2 ± 0.1 |

### Detection of residual zirconium released in the cell culture medium by the nanostructured pattern

The presence of traces of zirconium released by the nanostructured substrate into the culture medium has been evaluated by a spectrometric analysis. In line with the above-mentioned AFM data showing negligible variations of the roughness parameter R_q_ after incubation in the cell culture medium, the concentration of zirconium in the collected medium resulted to be lower than the limit of detection, i.e. <0.5 parts per billion (ppb) (Fig. S2). The geometric substrate partitioning fabricated with this procedure proved thus to be highly stable under cell culture conditions and to maintain their physical characteristics.

**Fig. S2.**
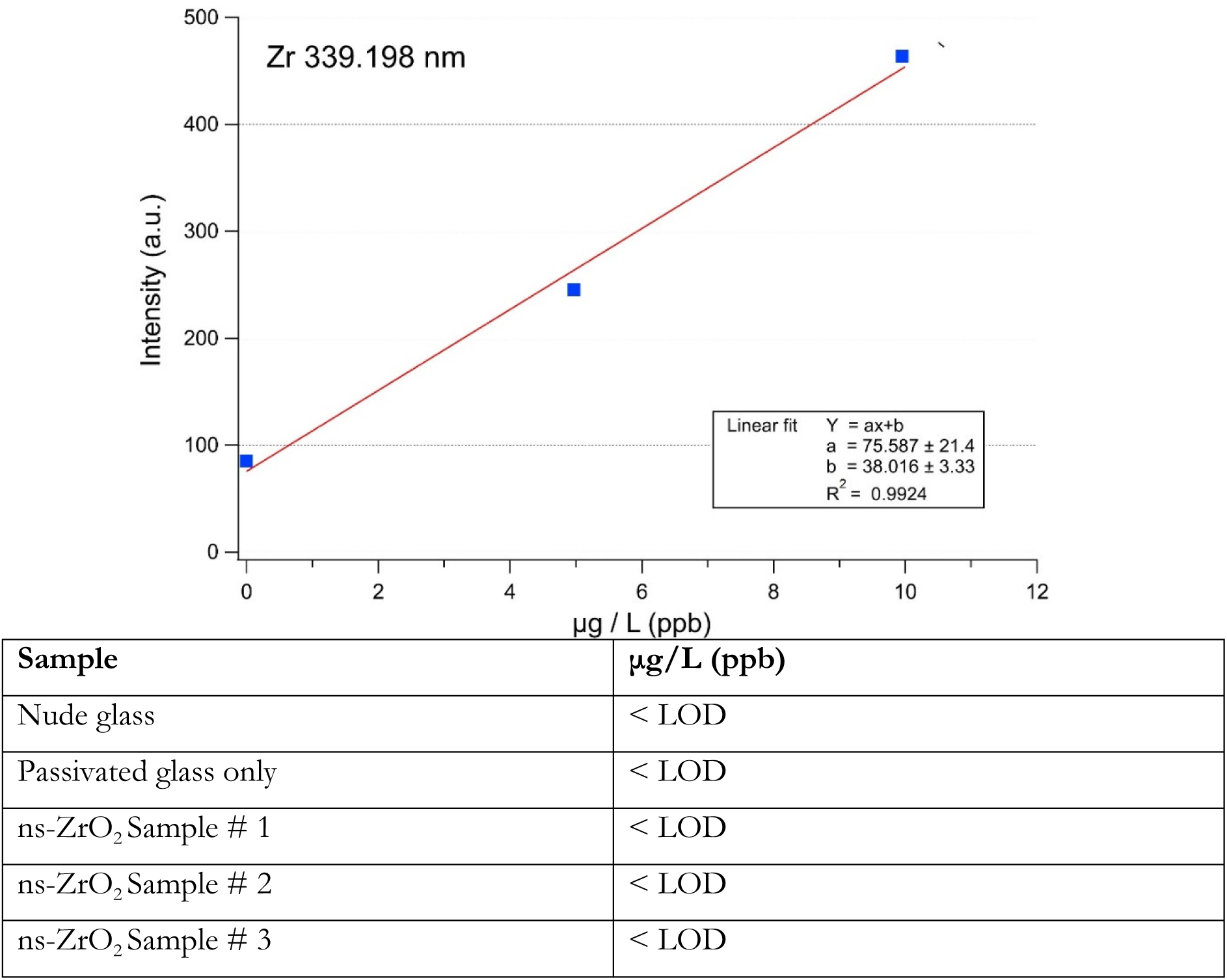
Top: Calibration curve of a spectrometric analysis obtained with the ICP-OES (iCAP 6200 Duo upgrade) for zirconium (Zr) (at the emission element wavelength of 339.198 nm); blank (= 0.0 ppb), standard 1 (= 4.970 ppb) and standard 2 (= 9.950 ppb) with 5LOD=2.5 ppb. Bottom: The table shows a quantitative analysis of the zirconium content for 2 negative controls (nude glass and passivated glass only) and 3 samples with micropatterned nanostructured zirconia substrates. For all the samples the concentration of zirconium results to be lower than the limit of detection, that is <0.5 ppb.

**Fig. S3.**
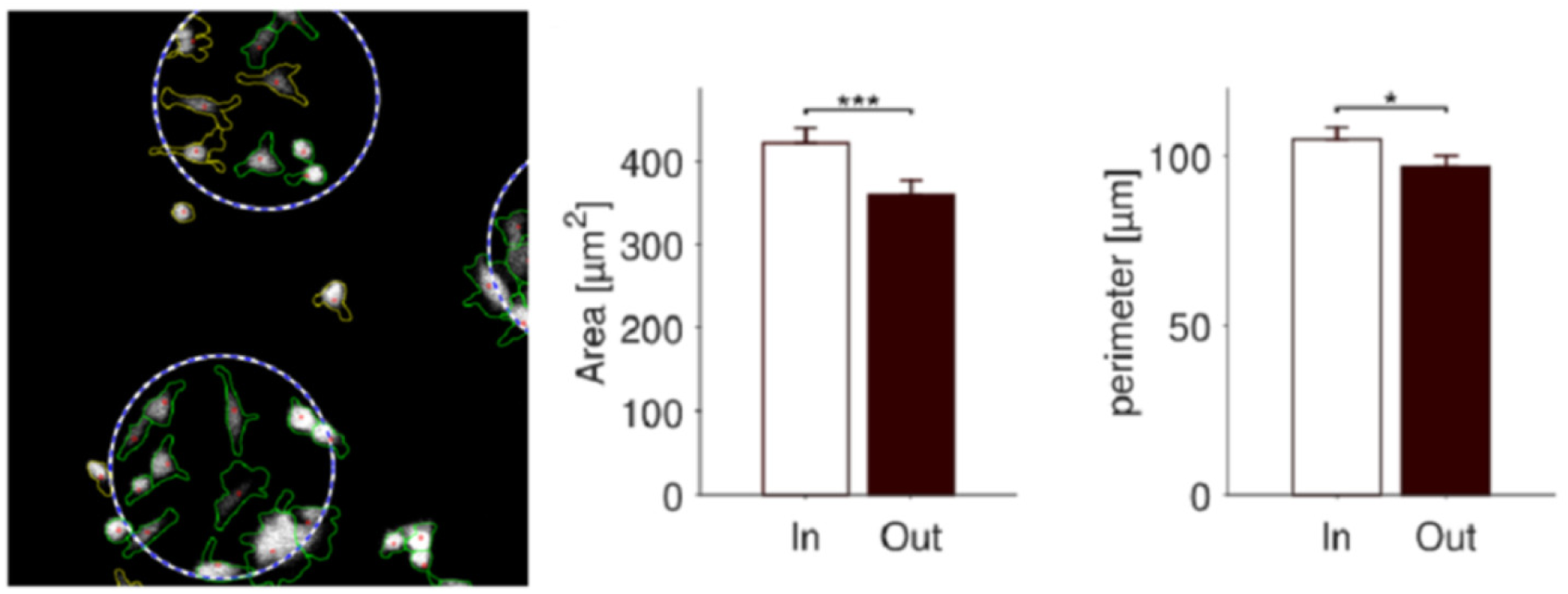
The image on the left shows an example of the cell shape recognition after application of the utilized algorithms(36)(37), and the graphs display the obtained values for the cell area (middle) and perimeter (right).

**Fig. S4.**
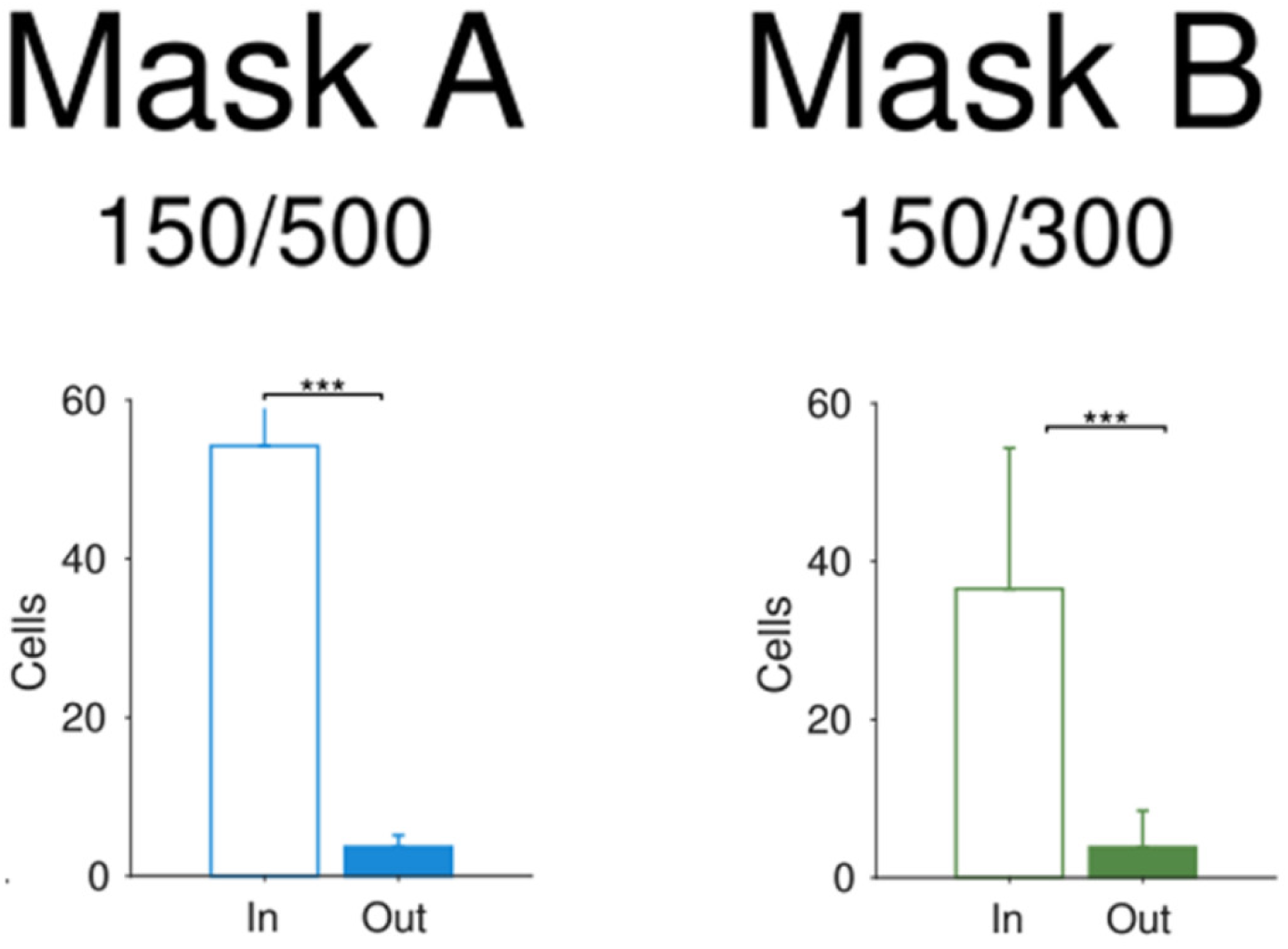
The average cell number on dot and outside dot, for the 50.000 cells/cm^2^ seeding concentration (see also Fig. 5).

**Fig. S5.**
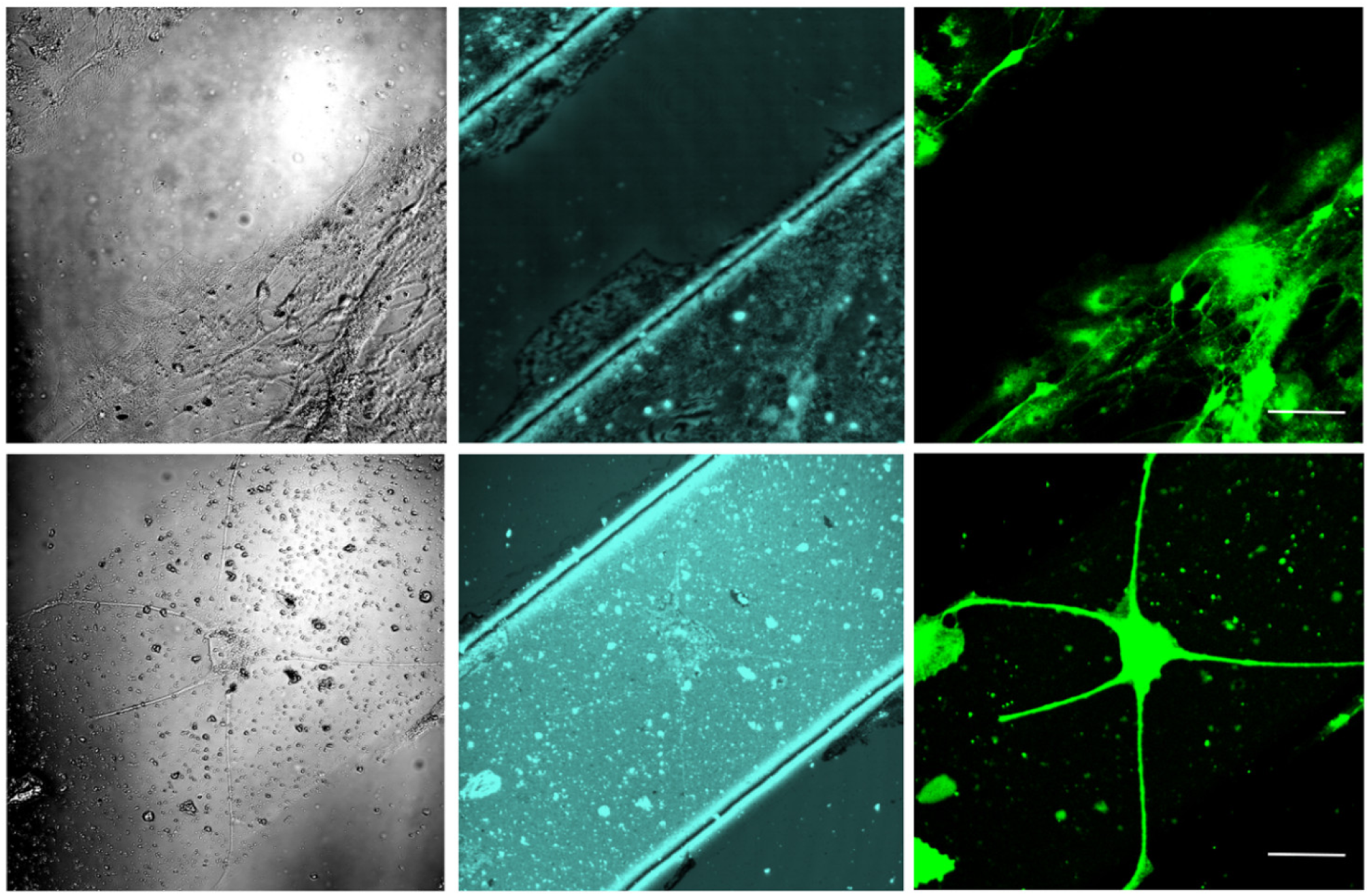
The panel shows further examples of primary hippocampal neonatal cells on a stripe-shaped pattern (which has not been further characterized for this work, but indicates the flexibility of the fabrication method). Left images: phase contrast, middle: confocal reflection, right: SynaptoZip expression (35), Scale bars = 50 μm.

